## Supplementary Materials for "Contributions of insula and superior temporal sulcus to interpersonal guilt and responsibility in social decisions"

**Supplementary Results**

**Supplementary Table 1: Mixed-effects regressions on choices**

For details of the models, see Results and Methods in the main text. *** p < 0.001; ** p < 0.01; * p < 0.05.

| | Study 1 probit | Study 1 linear | Study 2 probit | Study 2 linear |
|---|---|---|---|---|
| (Intercept) | 0.10 | 0.53*** | -0.04 | 0.49*** |
| | (0.08) | (0.02) | (0.07) | (0.02) |
| $EV_{risky} - V_{safe}$ | 0.07*** | 0.02*** | 0.09*** | 0.02*** |
| | (0.01) | (0.00) | (0.01) | (0.00) |
| Condition Social | 0.14* | 0.03^ | 0.01 | 0.01 |
| | (0.06) | (0.02) | (0.06) | (0.02) |
| $(EV_{risky} - V_{safe})$ * Condition | -0.00 | -0.00 | -0.00 | 0.00 |
| | (0.00) | (0.00) | (0.01) | (0.00) |
| $R^2$ (ord) | 1.000 | 0.284 | 1.000 | 0.306 |
| $R^2$ (adj) | 1.000 | 0.283 | 1.000 | 0.305 |
| AIC | 4909 | 5395 | 3864 | 4313 |
| BIC | 4974 | 5466 | 3927 | 4382 |
| LogLikelihood | -2445 | -2687 | -1922 | -2146 |
| N | 4680 | 4680 | 3833 | 3833 |

**Comparison between fits of the computational models to the momentary happiness data**

We compared the $R^2$ and adjusted $R^2$ values obtained for each participant and model using t-tests, for both studies. Average values across participants are shown in Table 1 of the main text. The *Responsibility* model yielded higher $R^2$ values than all the other models (Study 1: all $t > 3.6$, $p < 0.007$; Study 2: all $t > 2.9$, $p < 0.034$; Bonferroni-corrected t-tests) except for the *Guilt-envy* model in the data of Study 1 ($t = 2.19$, $p = 0.17$). The *Responsibility* and *Responsibility Redux* models yielded higher *adjusted* $R^2$ than the *Basic* model (Study 1: all $t > 3.6$, $p < 0.01$; Study 2: all $t > 3.6$, $p < 0.01$).

### Supplementary Figure 1: Parameter recovery for Responsibility Redux model.

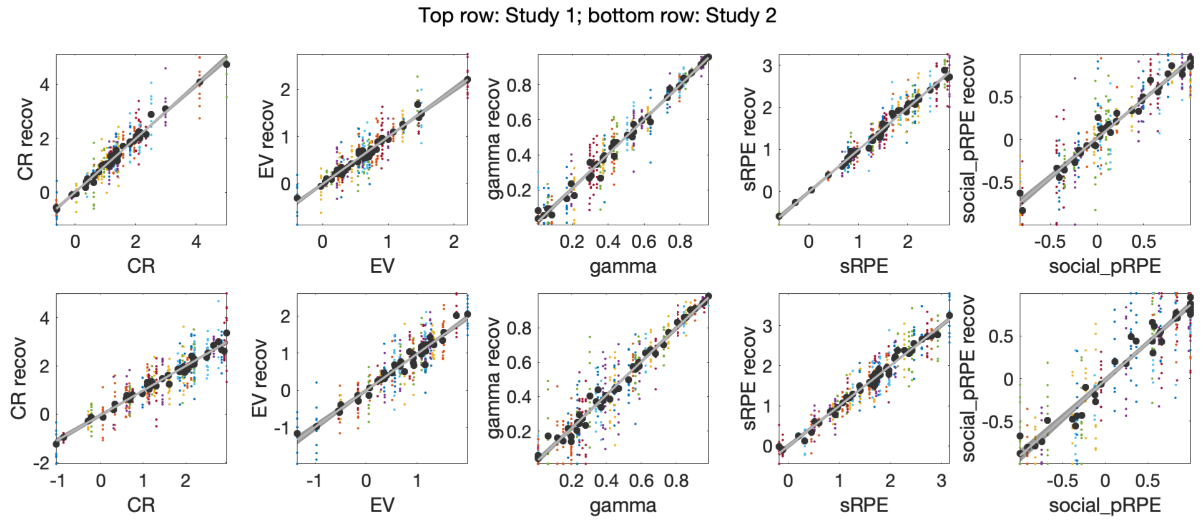


Stability of the estimated parameters of the temporal difference models was evaluated by attempting to recover parameters from synthetic data created using each participant's real estimated parameters. After fitting each participants' momentary happiness data (see above), we synthesized new momentary happiness data based on each participant's estimated parameters, added 1SD of noise to the happiness data, fitted the model to these synthetic data, and repeated this procedure 10 times, for both studies. We then compared these new estimated parameters to the actual parameters from which the synthetic data were generated. For each parameter, we calculated the mean of each participant's recovered parameters, and regressed these means on the participants' actual parameters (black dots and grey regression line, with 95% confidence interval). Each colored dot is one recovered parameter value; different colors represent different participants.

### Supplementary Table 2: linear mixed model regressions on happiness data following lottery choices

We fitted several models to the data in order to assess the stability of the effects. In both studies, Model 5 (Equation 9 in the Results section of the main text), which contained all 3 two-way interaction terms, explained the data best, so its parameters for the crucial partnerHigh:participantDecided interaction are reported in the main text. All models contained the three main fixed effects and subject as a random effect (random intercepts), models 2 through 4 contained one or two interaction terms. *** p < 0.001; ** p < 0.01; * p < 0.05.

**Study 1**

| | *Model 1* | *Model 2* | *Model 3* | *Model 4* | *Model 5* |
|---|---|---|---|---|---|
| (Intercept) | -0.58*** | -0.57*** | -0.47*** | -0.47*** | -0.38*** |
| | (0.06) | (0.06) | (0.06) | (0.07) | (0.07) |
| participantHigh [0,1] | 0.96*** | 0.95*** | 0.95*** | 0.94*** | 0.76*** |
| | (0.05) | (0.07) | (0.05) | (0.07) | (0.09) |
| partnerHigh [0,1] | 0.43*** | 0.43*** | 0.23** | 0.23** | 0.05 |
| | (0.05) | (0.05) | (0.07) | (0.07) | (0.09) |
| participantDecided [0,1] | -0.17** | -0.18* | -0.37*** | -0.39*** | -0.38*** |
| | (0.05) | (0.08) | (0.08) | (0.09) | (0.09) |
| participantHigh:participantDecided | - | 0.02 | - | 0.02 | 0.02 |
| | | (0.11) | | (0.11) | (0.1) |
| partnerHigh:participantDecided | - | - | 0.40*** | 0.40*** | 0.39*** |
| | | | (0.11) | (0.11) | (0.1) |
| participantHigh:partnerHigh | - | - | - | - | 0.35*** |
| | | | | | (0.1) |
| $R^2$ (ord) | 0.25 | 0.25 | 0.259 | 0.259 | 0.266 |
| $R^2$ (adj) | 0.248 | 0.247 | 0.256 | 0.256 | 0.262 |
| N | 1216 | 1216 | 1216 | 1216 | 1216 |

**Study 2**

| | *Model 1* | *Model 2* | *Model 3* | *Model 4* | *Model 5* |
|---|---|---|---|---|---|
| (Intercept) | -0.54*** | -0.50*** | -0.45*** | -0.43*** | -0.26** |
| | (0.06) | (0.07) | (0.07) | (0.08) | (0.09) |
| participantHigh [0,1] | 1.01*** | 0.95*** | 1.02*** | 0.96*** | 0.65*** |
| | (0.06) | (0.09) | (0.06) | (0.09) | (0.11) |
| partnerHigh [0,1] | 0.35*** | 0.35*** | 0.20* | 0.20* | -0.11 |
| | (0.06) | (0.06) | (0.09) | (0.09) | (0.11) |
| participantDecided [0,1] | -0.22*** | -0.28** | -0.38*** | -0.43*** | -0.45*** |
| | (0.06) | (0.09) | (0.09) | (0.11) | (0.11) |
| participantHigh:participantDec. | - | 0.11 | - | 0.1 | 0.12 |
| | | (0.12) | | (0.12) | (0.12) |
| partnerHigh:participantDecided | - | - | 0.29* | 0.28* | 0.31** |
| | | | (0.12) | (0.12) | (0.12) |
| participantHigh:partnerHigh | - | - | - | - | 0.58*** |
| | | | | | (0.12) |
| $R^2$ (ord) | 0.28 | 0.281 | 0.285 | 0.285 | 0.304 |
| $R^2$ (adj) | 0.278 | 0.278 | 0.282 | 0.281 | 0.299 |
| N | 876 | 876 | 876 | 876 | 876 |

**Supplementary Table 3: Details of clusters with activation varying as a function of choice and condition, during the decision phase (output of second-level SPM model plus Cohen's d for each cluster)**

| *Anatomy* | *Size (N vox.)* | *T* | *d* | *Z* | *MNI* | | |
|---|---|---|---|---|---|---|---|
| | | | | | *x* | *y* | *z* |
| **Risky > safe** | | | | | | | |
| VStriatum R | 385 | 7.14 | 0.79 | 6.78 | 10 | 12 | -4 |
| VStriatum R | - | 5.28 | - | 5.12 | 14 | 22 | 4 |
| VStriatum L | 393 | 6.15 | 0.62 | 5.92 | -10 | 8 | -6 |
| **Social > solo** | | | | | | | |
| Precuneus R | 737 | 5.72 | 0.63 | 5.53 | 0 | -62 | 38 |
| Precuneus L | - | 5.17 | - | 5.02 | -10 | -56 | 34 |
| Precuneus L | - | 4.91 | - | 4.79 | -2 | -54 | 34 |
| Angular L (TPJ) | 320 | 4.50 | 0.55 | 4.40 | -34 | -58 | 26 |
| Temporal Sup L | - | 3.59 | - | 3.54 | -52 | -56 | 20 |
| Medial PFC R | 302 | 3.97 | 0.54 | 3.90 | 4 | 52 | 22 |
| Medial PFC L | - | 3.97 | - | 3.90 | -6 | 58 | 22 |
| Medial PFC L | - | 3.63 | - | 3.57 | -6 | 54 | 32 |

**Supplementary Table 4: Linear mixed model regressions on BOLD response parameter estimates obtained during the decision phase**

a) **Analysis of the response during choice**

The parameter estimates of all voxels of the ROIs identified using the contrasts Risky > Safe and Social > Solo (see above and main text) were fitted with linear mixed models. The parameters of the best-fitting model (lowest BIC) for each ROI are reported below. We note here that the *Run* factor and interactions with it had significant effects in several ROIs, which shows that some of the effects reported varied across runs. However, for the sake of brevity, we will not discuss these results further. *** p < 0.001; ** p < 0.01; * p < 0.05.

| | *VStria L* | *VStria R* | *MPFC* | *Precun* | *TPJ L* |
|---|---|---|---|---|---|
| (Intercept) | -0.78*** | -1.52*** | 1.18*** | 1.39*** | 0.94*** |
| | (0.13) | (0.13) | (0.18) | (0.18) | (0.14) |
| choice | -0.26*** | 0.78*** | -0.38*** | 0.97*** | 0.55*** |
| | (0.08) | (0.07) | (0.09) | (0.07) | (0.08) |
| condition | 0.01 | 0.34*** | -0.63*** | -0.40*** | -0.26*** |
| | (0.03) | (0.02) | (0.03) | (0.02) | (0.03) |

| | | | | | |
|---|---|---|---|---|---|
| run | 0.32*** | 0.96*** | -0.26*** | 0.30*** | 0.09* |
| | (0.04) | (0.03) | (0.04) | (0.03) | (0.04) |
| choice:condition | 0.47*** | 0.04 | 0.12** | -0.30*** | -0.24*** |
| | (0.04) | (0.03) | (0.04) | (0.03) | (0.04) |
| choice:run | 0.33*** | -0.29*** | 0.09 | -0.56*** | -0.37*** |
| | (0.05) | (0.05) | (0.06) | (0.05) | (0.05) |
| condition:run | -0.01 | -0.22*** | 0.17*** | -0.02 | -0.02 |
| | (0.02) | (0.02) | (0.02) | (0.01) | (0.02) |
| choice:condition:run | -0.18*** | 0.05* | -0.01 | 0.16*** | 0.12*** |
| | (0.02) | (0.02) | (0.03) | (0.02) | (0.02) |
| $R^2$ (ord) | 0.136 | 0.158 | 0.208 | 0.170 | 0.175 |
| $R^2$ (adj) | 0.136 | 0.158 | 0.208 | 0.170 | 0.175 |

**b) Analysis of the difference between Risky and Safe choices, Social vs Solo**

To better understand the choice:condition interaction, which was significant in all ROIs except the right striatum, we subtracted the response to safe choices from the response to risky choices for the four remaining ROIs and submitted these differences to additional linear mixed models, as above. The first model contained a factor socialVsSolo, in which data from the social condition were weighted positively, and trials in the solo condition were weighted negatively. As above, we tested these models both with and without the factor *Run* and associated interaction, and we report the best-fitting model in the table below: a dash ('-') in the row displaying parameters for the *run* and *socialVsSolo:run* regressors indicates that the model without factor *run* was better-fitting for this ROI.

| | *VStria L* | *MPFC* | *Precun* | *TPJ L* |
|---|---|---|---|---|
| (Intercept) | 0.67*** | -0.03 | 0.38*** | 0.07 |
| | (0.11) | (0.10) | (0.11) | (0.09) |
| socialVsSolo | -0.47*** | -0.10*** | 0.30*** | 0.24*** |
| | (0.03) | (0.01) | (0.02) | (0.03) |
| run | -0.03* | - | -0.24*** | -0.14*** |
| | (0.02) | | (0.01) | (0.01) |
| socialVsSolo:run | 0.18*** | - | -0.16*** | -0.12*** |
| | (0.02) | | (0.01) | (0.02) |
| R2 (ord) | 0.085 | 0.075 | 0.068 | 0.091 |
| R2 (adj) | 0.085 | 0.075 | 0.067 | 0.091 |
| N | 94320 | 72480 | 176880 | 76800 |

c) **Analysis of the difference between Risky and Safe choices, Social vs Partner**

Finally, we repeated this analysis with models containing a factor socialVsPartner, in which data from the social condition were weighted positively, and trials in the partner condition were weighted negatively. Here again, we report the best-fitting model from the versions with and without the factor *run*.

| | *VStria L* | *MPFC* | *Precun* | *TPJ L* |
|---|---|---|---|---|
| (Intercept) | 0.61*** | -0.03 | 0.38*** | 0.07 |
| | (0.11) | (0.10) | (0.11) | (0.09) |
| socialVsPartner | -0.07*** | 0.23*** | 0.39*** | 0.07** |
| | (0.01) | (0.01) | (0.02) | (0.03) |
| run | - | - | -0.24*** | -0.14*** |
| | | | (0.01) | (0.01) |
| socialVsPartner:run | - | - | -0.20*** | 0.01 |
| | | | (0.01) | (0.02) |
| $R^2$ (ord) | 0.090 | 0.225 | 0.233 | 0.228 |
| $R^2$ (adj) | 0.090 | 0.225 | 0.233 | 0.228 |
| N | 94320 | 72480 | 176880 | 76800 |

**Supplementary Table 5: Details of clusters with higher activation during risky vs. safe outcomes (second-level SPM model, with Cohen's d for each cluster).**

Note: A dash ('-') in the *Size* or *d* column indicates that the peak reported on that line is part of a cluster whose centre is the next peak without dash listed above it.

| *Anatomy* | *Size (N v.)* | *T* | *d* | *Z* | *x* | *y* | *z* |
|---|---|---|---|---|---|---|---|
| Insula R | 1586 | 11.87 | 2.27 | Inf | 30 | 22 | -10 |
| | - | 10.51 | - | Inf | 42 | 22 | -8 |
| | - | 7.11 | - | 6.99 | 50 | 22 | 6 |
| Insula L | 1164 | 11.8 | 1.98 | Inf | -30 | 20 | -10 |
| | - | 6.15 | - | 6.07 | -50 | 16 | 8 |
| Dorsomedial_PFC | 2962 | 9.70 | 2.38 | Inf | 2 | 42 | 36 |
| | - | 8.31 | - | Inf | 2 | 20 | 60 |
| | - | 7.99 | - | 7.81 | 4 | 40 | 20 |
| Temp_Mid_R (STS) | 80 | 6.06 | 1.14 | 5.98 | 48 | -24 | -8 |
| | - | 4.93 | - | 4.89 | 50 | -34 | -2 |
| VStria_L | 30 | 5.73 | 1.23 | 5.66 | -10 | 0 | -6 |

| | | | | | | | |
|---|---|---|---|---|---|---|---|
| VStria_R | 40 | 5.68 | 1.18 | 5.62 | 8 | 4 | 2 |
| Parietal_Inf_R | 193 | 5.64 | 1.55 | 5.58 | 40 | -48 | 46 |
| | - | 5.12 | - | 5.07 | 36 | -60 | 54 |
| | - | 5.09 | - | 5.05 | 48 | -36 | 48 |
| DSL_PFC_R | 56 | 5.51 | 1.38 | 5.46 | 42 | 38 | 24 |
| Parietal_Inf_L | 64 | 5.27 | 1.25 | 5.21 | -46 | -44 | 46 |
| | - | 5.03 | - | 4.99 | -38 | -42 | 38 |

**Supplementary Table 6: Linear mixed model regressions on BOLD response parameter estimates obtained during the outcome phase**

The parameter estimates of all voxels of the nine ROIs identified using the contrasts Risky > Safe outcome (see main Text) were fitted with linear mixed models. The parameters of the best-fitting model (lowest BIC) for each ROI are reported below. *Social* was a dummy variable with the value of 1 for the *Social* condition and 0 for the *Partner* condition; *LowOutcome* was a dummy with the value of 1 or *Low lottery outcome* and 0 for *High lottery outcome*; *Run* was a dummy with the value of 1 for run 1 and 2 for run 2. Subject was the only random factor. We note here that the *Run* factor and interactions with it were significant in several ROIs, which indicates that some of the effects reported varied across runs. However, for the sake of brevity, we will not discuss these results further. *** $p < 0.001$; ** $p < 0.01$; * $p < 0.05$; ^ $p < 0.1$.

| | **VStriaL** | **VStriaR** | **InsulaL** | **InsulaR** | **STS_R** |
|---|---|---|---|---|---|
| (Intercept) | 1.79*** | 1.12*** | 0.73*** | 0.52*** | 1.09*** |
| | (0.25) | (0.27) | (0.14) | (0.15) | (0.18) |
| Social | -1.30*** | -0.68*** | 0.42*** | 0.42*** | 0.21^ |
| | (0.22) | (0.08) | (0.04) | (0.03) | (0.12) |
| LowOutcome | -1.14*** | -0.12 | 0.83*** | 0.58*** | 0.51*** |
| | (0.22) | (0.08) | (0.04) | (0.03) | (0.12) |
| Run | -0.82*** | - | 0.18*** | 0.34*** | 0.40*** |
| | (0.10) | | (0.02) | (0.02) | (0.06) |
| Social:LowOutcome | 0.95** | 0.77*** | -0.76*** | -0.25*** | -0.81*** |
| | (0.31) | (0.12) | (0.06) | (0.05) | (0.18) |
| Social:Run | 0.89*** | - | -0.30*** | -0.29*** | -0.67*** |
| | (0.14) | | (0.03) | (0.02) | (0.08) |
| LowOutcome:Run | 0.60*** | - | -0.75*** | -0.43*** | -0.66*** |
| | (0.14) | | (0.03) | (0.02) | (0.08) |
| Social:LowOutcome:Run | -0.30 | - | 0.80*** | 0.30*** | 0.99*** |
| | (0.20) | | (0.04) | (0.03) | (0.11) |
| R2 (ord) | 0.200 | 0.201 | 0.086 | 0.099 | 0.170 |

| | | | | | |
|---|---|---|---|---|---|
| R2 (adj) | 0.200 | 0.201 | 0.086 | 0.099 | 0.170 |
| N | 9600 | 12800 | 372480 | 507520 | 25600 |

| | **ParL** | **ParR** | **dmPFC** | **FrontR** |
|---|---|---|---|---|
| (Intercept) | -0.22 | 0.96*** | 0.73*** | -0.22 |
| | (0.21) | (0.22) | (0.10) | (0.21) |
| Social | 0.14 | -0.60*** | -0.48*** | -0.07 |
| | (0.13) | (0.09) | (0.02) | (0.15) |
| LowOutcome | 0.74*** | -0.81*** | 0.16*** | -0.80*** |
| | (0.13) | (0.09) | (0.02) | (0.15) |
| Run | 0.94*** | 0.69*** | 0.24*** | 0.90*** |
| | (0.06) | (0.04) | (0.01) | (0.07) |
| Social:LowOutcome | 0.12 | 2.17*** | 0.69*** | 2.05*** |
| | (0.18) | (0.13) | (0.03) | (0.22) |
| Social:Run | -0.29*** | 0.00 | 0.18*** | -0.23* |
| | (0.08) | (0.06) | (0.01) | (0.10) |
| LowOutcome:Run | -0.54*** | 0.41*** | -0.17*** | 0.53*** |
| | (0.08) | (0.06) | (0.01) | (0.10) |
| Social:LowOutcome:Run | 0.26* | -0.88*** | -0.33*** | -1.21*** |
| | (0.12) | (0.08) | (0.02) | (0.14) |
| $R^2$ (ord) | 0.255 | 0.220 | 0.063 | 0.218 |
| $R^2$ (adj) | 0.255 | 0.220 | 0.063 | 0.218 |
| N | 20480 | 61760 | 947840 | 17920 |

#### a) Analysis of the response during low lottery outcomes

To identify regions likely to be involved in the guilt effect, we focused on the regions engaged when participants rather than their partner made the choice, i.e. the regions responding significantly more to the *Social* than the *Partner* condition. Of these regions, the insulae and the right middle temporal cortex also showed a significant *Social:LowOutcome* interaction. To better understand this interaction in these three ROIs, we ran additional models to test the effect of the *Social* compared to the *Partner* condition on the responses to *Low lottery outcomes* only, and on their response difference between *Low* and *High lottery outcomes*. The results for the response to low lottery outcomes were:

| | **InsulaL** | **InsulaR** | **MidTempR** |
|---|---|---|---|
| (Intercept) | 0.70*** | 0.95*** | 1.22*** |
| | (0.18) | (0.18) | (0.18) |
| Social | 0.41*** | 0.18*** | -0.12** |
| | (0.01) | (0.01) | (0.04) |
| $R^2$ (ord) | 0.150 | 0.148 | 0.231 |
| $R^2$ (adj) | 0.150 | 0.148 | 0.231 |

| | | | |
|---|---|---|---|
| N | 186240 | 253760 | 12800 |

b) **Analysis of the response during low lottery outcomes**

The results of the models fitted to the response difference (response to *Low lottery outcomes* minus response to *High lottery outcomes*) were:

| | **InsulaL** | **InsulaR** | **MidTempR** |
|---|---|---|---|
| (Intercept) | -0.30 | -0.07 | -0.48* |
| | (0.19) | (0.16) | (0.23) |
| Social | 0.44*** | 0.19*** | 0.67*** |
| | (0.02) | (0.01) | (0.04) |
| $R^2$ (ord) | 0.116 | 0.091 | 0.253 |
| $R^2$ (adj) | 0.116 | 0.091 | 0.253 |
| N | 186240 | 253760 | 12800 |

Of all these regions, only the left and right insulae showed a higher response to the *Social* compared to the *Partner* condition, a significant interaction between *Social* vs *Partner* and *Low* vs *High lottery outcome*, and higher responses to *Low lottery outcomes* when these resulted from participants' choices.

**Supplementary Figure 2: Functional connectivity with the left TD-model-defined STS (seed) during choices in *Solo* and *Social* conditions.**

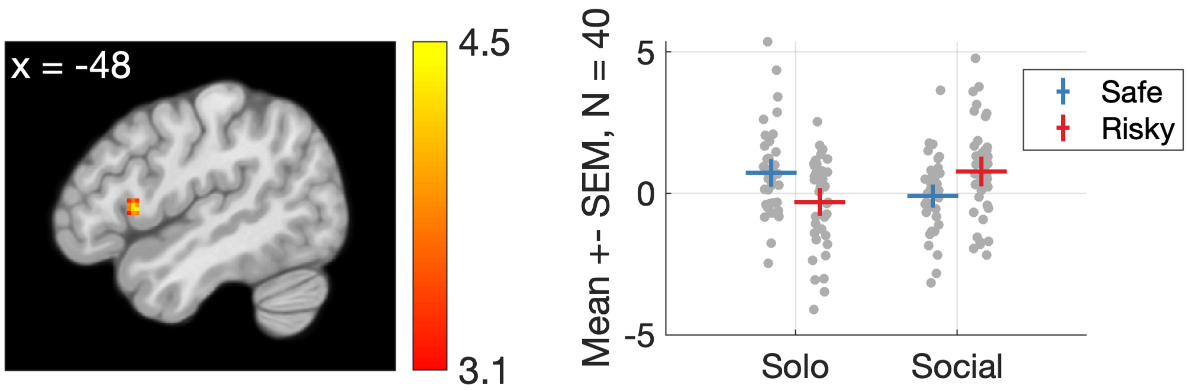


*Connectivity between the left STS (seed) and a cluster in the left inferior frontal gyrus that did not survive corrections for multiple tests, where connectivity with the left STS (the seed region) showed the opposite pattern: connectivity was highest when participants made Safe choices for themselves and Risky choices for both players ($p_{uncorrected}$ = 0.001, T=4.44, Z=4.30, 35 voxels, peak at MNI [-48 14 6]. In the*

*righthand panel, dots are data of individual participants, the markers represent means, and error bars indicate 95 confidence intervals about the mean.*

**Supplementary Table 7: Participant's judgments of their partner**

| | *Study 1* | *Study 2* |
|---|---|---|
| How sympathetic did you find them? | 8.56 (1.7) | 9.18 (1.17) |
| How well could you cooperate with them? | 8.35 (1.29) | 8.68 (1.18) |
| How honest did they seem? | 9.05 (1.11) | 9.34 (1.10) |
| How open were they? | 8.43 (1.89) | 9.05 (1.12) |
| How sociable were they? | 8.63 (1.61) | 9.28 (1.18) |

Scale used was 1 (minimum) to 10 (maximum). Mean and standard deviations are reported.
